# Networks extracted from nonlinear fMRI connectivity exhibit unique spatial variation and enhanced sensitivity to differences between individuals with schizophrenia and controls

**DOI:** 10.1101/2023.11.16.566292

**Authors:** Spencer Kinsey, Katarzyna Kazimierczak, Pablo Andrés Camazón, Jiayu Chen, Tülay Adali, Peter Kochunov, Bhim Adhikari, Judith Ford, Theo G. M. van Erp, Mukesh Dhamala, Vince D. Calhoun, Armin Iraji

## Abstract

Functional magnetic resonance imaging (fMRI) studies often estimate brain intrinsic connectivity networks (ICNs) from temporal relationships between hemodynamic signals using approaches such as independent component analysis (ICA). While ICNs are thought to represent functional sources that play important roles in various psychological phenomena, current approaches have been tailored to identify ICNs that mainly reflect linear statistical relationships. However, the elements comprising neural systems often exhibit remarkably complex nonlinear interactions that may be involved in cognitive operations and altered in psychiatric conditions such as schizophrenia. Consequently, there is a need to develop methods capable of effectively capturing ICNs from measures that are sensitive to nonlinear relationships. Here, we advance a novel approach to estimate ICNs from explicitly nonlinear whole-brain functional connectivity (ENL-wFC) by transforming resting-state fMRI (rsfMRI) data into the connectivity domain, allowing us to capture unique information from distance correlation patterns that would be missed by linear whole-brain functional connectivity (LIN-wFC) analysis.

Our findings provide evidence that ICNs commonly extracted from linear (LIN) relationships are also reflected in explicitly nonlinear (ENL) connectivity patterns. ENL ICN estimates exhibit higher reliability and stability, highlighting our approach’s ability to effectively quantify ICNs from rsfMRI data. Additionally, we observed a consistent spatial gradient pattern between LIN and ENL ICNs with higher ENL weight in core ICN regions, suggesting that ICN function may be subserved by nonlinear processes concentrated within network centers. We also found that a uniquely identified ENL ICN distinguished individuals with schizophrenia from healthy controls while a uniquely identified LIN ICN did not, emphasizing the valuable complementary information that can be gained by incorporating measures that are sensitive to nonlinearity in future analyses. Moreover, the ENL estimates of ICNs associated with auditory, linguistic, sensorimotor, and self-referential processes exhibit heightened sensitivity towards differentiating between individuals with schizophrenia and controls compared to LIN counterparts, demonstrating the translational value of our approach and of the ENL estimates of ICNs that are frequently reported as disrupted in schizophrenia. In summary, our findings underscore the tremendous potential of connectivity domain ICA and nonlinear information in resolving complex brain phenomena and revolutionizing the landscape of clinical FC analysis.

## 1. Introduction

Brain function is underpinned by interacting assemblies of neurons organized at various spatial and temporal scales. At the whole-brain level, functional magnetic resonance imaging (fMRI) analysis is a non-invasive tool that has commonly been used to estimate coherent neuronal ensembles from statistical relationships between blood-oxygenation-level-dependent (BOLD) time series, which is often called functional connectivity (FC). Although the relationship between the BOLD signal and neural activity is indirect (Logothetis et al., 2003), experimentally induced and resting-state BOLD fluctuations are typically associated with changes in local field potentials across multiple frequency bands (Logothetis et al., 2001; Magri et al., 2012; Pan et al., 2013; Shi et al., 2017), indicating that fMRI FC analysis is a promising tool for advancing the identification of task-related and spontaneously emerging networks of interacting brain regions. Moreover, fMRI FC analysis is deployable within a wide range of predictive clinical contexts. For example, multiple large-scale meta-analyses have shown that FC can be used to distinguish healthy controls (HC) from individuals with schizophrenia (SZ) (Dong et al., 2018; Li et al., 2019; Sheffield & Barch, 2016). Such studies have contributed to an accumulation of evidence for the SZ “dysconnection hypothesis” (Friston et al., 2016), which posits FC alteration as a central endophenotype of the disorder resulting from neuromodulatory and synaptic pathogenesis.

However, FC studies are typically designed to estimate networks that reflect linear statistical relationships between brain areas (Friston, 2011; Mohanty et al., 2020). As a result, the remarkably complex nonlinear interactions inherent to neural systems have remained largely outside of the focal lens of FC research (Friston, 2001; He & Yang, 2021; Singer, 2013). We highlight three ways in which bringing nonlinear dependence to the fore has potential to advance systems, cognitive, and predictive neuroscientific research. First, the analysis of nonlinear statistical relationships may lead to a more precise and thorough characterization of the organization and dynamics of neural ensembles at multiple scales (He & Yang, 2021; Iraji et al., 2022b; Iraji et al., 2023a). Second, nonlinear interactions may have cognitive and behavioral relevance. In principle, nonlinearity is thought to underpin a high-dimensional state space capable of supporting a set of flexible and diverse neural computations (Friston, 2001; Singer, 2013), such that analyzing its functional role may shed light on the structure of cognitive processing and the deficiencies associated with psychiatric disorders and symptoms. Third, measures that are sensitive to nonlinearity can be leveraged to develop biomarkers that can be incorporated within brain-based predictive models of mental illness, or “predictomes” (Rashid & Calhoun, 2020).

Among available FC analysis methods, independent component analysis (ICA) is a powerful multivariate source separation technique that has been applied to fMRI data. ICA assumes that the data is a linear mixture of statistically independent source signals and aims to estimate an unmixing matrix yielding components that optimally approximate these signals (Adali et al., 2014; Comon & Jutten, 2010). In the context of fMRI analysis, spatial ICA (sICA) has commonly been used to decompose fMRI time series data into a set of intrinsic connectivity networks (ICNs), where the spatial pattern of an ICN describes its distribution across voxels and the temporal pattern describes its activity over time (Calhoun et al., 2008; Calhoun et al., 2009; Iraji et al., 2022a; Seeley et al., 2007). ICNs can be robustly and consistently identified from both resting-state fMRI (rsfMRI) (Calhoun et al., 2008; Damoiseaux et al., 2006; Iraji et al., 2022a) and task-based fMRI (tfMRI) time series data (Calhoun et al., 2008; Calhoun et al., 2009; Laird et al., 2011; Wu et al., 2021) at different spatial scales (Iraji et al., 2019a; Iraji et al., 2022b; Iraji et al., 2023a). ICNs can also be reliably extracted from FC matrices constructed from second-order statistics such as Pearson correlation (i.e., from the connectivity domain) (Iraji et al., 2016; Wu et al., 2018). Connectivity domain ICA is a type of feature-based analysis (Calhoun & Allen, 2013) that yields cross-validating components and is distinguished from time domain ICA by unique benefits including consistency across changes in particular analysis parameters and reproducibility (Iraji et al., 2016). Moreover, the Pearson correlation coefficient is one out of an expanding range of FC metrics that can be used to construct the connectivity basis, making connectivity domain ICA an incredibly versatile tool (Iraji et al., 2016).

Connectivity domain and time domain ICA have become valuable tools for investigating fMRI data. However, both methods are typically designed to identify ICNs comprised of covarying brain regions, reflecting linear FC (Calhoun et al., 2009; Iraji et al., 2016; Wu et al., 2018). Although recent advancements have made strides in incorporating nonlinearity (Hyvärinen et al., 2019; Morioka et al., 2020) such as learning local spatial or temporal nonlinear structures, the estimated functional sources still do not effectively quantify nonlinear FC. Here, we advance a novel connectivity domain ICA approach to extract ICNs from explicitly nonlinear whole-brain FC (ENL-wFC) estimated from distance correlation (Székely et al., 2007) patterns that move beyond those constructed from Pearson correlation. Although alternative metrics can be used to quantify fMRI connectivity while accounting for higher-order statistics (Bhinge et al., 2019; Motlaghian et al., 2022; Motlaghian et al., 2023), distance correlation is a powerful and flexible dependence metric that remains relatively underexplored in the context of FC research. Moreover, the proposed method is unique in that we conceive of ENL-wFC as a global feature of the connectivity space, rather than as a composite feature constructed from the explicitly nonlinear associations between time series pairs (Motlaghian et al., 2022; Motlaghian et al., 2023). This allows us to leverage the potentially valuable information present within global connectivity features that move beyond the connectivity features constructed from Pearson correlation.

We find that our method is capable of effectively extracting ICNs from ENL-wFC. Our results show that many ICNs extracted from linear whole-brain FC (LIN-wFC) information are also reflected within ENL-wFC, although we find that ENL ICNs are estimated more reliably, and that corresponding ICNs exhibit graded spatial variation with greater ENL weight within many core regions. We also identify unique ENL and LIN ICNs, indicating that our approach can recover ICNs that are hidden from conventional linear FC analyses. Furthermore, we find that ENL ICN estimates are more sensitive to differences between HC and SZ overall in addition to being more sensitive for specific ICNs that have been previously reported as altered in SZ, demonstrating that ENL ICNs can be leveraged within a predictive clinical context. More generally, we demonstrate that connectivity domain ICA is a powerful and flexible tool that is uniquely poised to transform clinical FC analysis and advance our understanding of brain function.

## 2. Materials and Methods

### 2.1 Subject Information, Data Acquisition, and Subject Quality Control Criteria

We analyzed 3-Tesla resting-state fMRI (rsfMRI) time series data sourced from three psychosis projects: Center for Biomedical Research Excellence (COBRE), Functional Imaging Biomedical Informatics Research Network (FBIRN), and Maryland Psychiatric Research Center (MPRC) (**Table 1**). Detailed subject recruitment information for COBRE, FBIRN, and MPRC studies can be respectively found in Aine et al. (2017), Damaraju et al. (2014), and Adhikari et al. (2019).

**Table 1.** Subject demographic information. COBRE: Center for Biomedical Research Excellence. FBIRN: Functional Imaging Biomedical Informatics Research Network. MPRC: Maryland Psychiatric Research Center. HC: healthy control. SZ: schizophrenia.

| Dataset | Diagnosis (#) | Sex (#) | Age (years) mean ± sd | Median / range |
| --- | --- | --- | --- | --- |
| COBRE | HC (75) | Male (56) | 39.27 ± 12.13 | 39 / (65 – 18) |
| | | Female (19) | 35.47 ± 10.02 | 34 / (58 – 18) |
| | SZ (51) | Male (45) | 37.36 ± 15.28 | 33 / (64 – 19) |
| | | Female (6) | 40.83 ± 17.70 | 44 / (65 – 20) |
| FBIRN | HC (88) | Male (60) | 36.58 ± 10.74 | 39 / (59 – 19) |
| | | Female (28) | 36.61 ± 11.07 | 33 / (58 – 19) |
| | SZ (60) | Male (52) | 39.54 ± 11.12 | 41 / (60 – 18) |
| | | Female (8) | 35.88 ± 9.85 | 34 / (51 – 24) |
| MPRC | HC (152) | Male (69) | 38.99 ± 13.22 | 41 / (68 – 18) |
| | | Female (83) | 39.93 ± 14.93 | 43 / (64 – 16) |
| | SZ (82) | Male (57) | 36.25 ± 13.54 | 33 / (63 – 13) |
| | | Female (25) | 44.68 ± 11.92 | 47 / (61 – 13) |

Individuals with SZ from the COBRE dataset received a diagnosis of schizophrenia performed in consensus by two research psychiatrists via the Structured Clinical Interview for DSM-IV Axis I Disorders (SCID) using the patient version of the SCID-DSM-IV-TR (Aine et al., 2017). SZ subjects were evaluated for comorbidities and for retrospective as well as prospective clinical stability. Individuals with SZ from the FBIRN study were diagnosed with schizophrenia based on the SCID-DSM-IV-TR and were clinically stable for at least two months prior to scanning (Damaraju et al., 2014). For MPRC SZ subjects, a diagnosis of schizophrenia was confirmed via the SCID-DSM-IV (Adhikari et al., 2019).

COBRE data were collected at a single site on a Siemens TIM Trio scanner via an echo-planar imaging sequence (TR = 2000 ms; TE = 29 ms) (Iraji et al., 2022b). Voxel spacing was 3.75 x 3.75 x 4.5 mm, the slice gap was 1.05 mm, and the field of view (FOV) was 240 x 240 mm. FBIRN data were collected from seven sites (Turner et al., 2013), with six sites utilizing Siemens TIM Trio scanners and one utilizing a General Electric Discovery MR750 (Iraji et al., 2022b). All seven sites used an echo-planar imaging sequence (TR = 2000 ms; TE = 30 ms). Original voxel spacing was 3.4375 x 3.4375 x 4 mm, the slice gap was 1 mm, and the FOV was 220 x 220 mm. MPRC data were collected from three sites via echo-planar imaging sequences (Friston, 2011; Iraji et al., 2022b). One site used a Siemens Allegra scanner (TR = 2000 ms; TE = 27 ms; voxel spacing = 3.44 x 3.44 x 4 mm; FOV = 220 x 220 mm), another used a Siemens TIM Trio scanner (TR = 2210 ms; TE = 30 ms; voxel spacing = 3.44 x 3.44 x 4 mm; FOV = 220 x 220 mm), and the third site used a Siemens TIM Trio scanner (TR = 2000 ms; TE = 30 ms; voxel spacing = 1.72 x 1.72 x 4 mm; FOV = 220 x 220 mm).

The following subject quality control criteria (Iraji et al., 2023a) were used for the current study: 1) maximum head rotations of less than 3⁰ and maximum translations of less than 3 mm, 2) mean framewise displacement (FD) less than 0.25, 3) quality registration to an echo-planar imaging template, 4) and whole-brain (in addition to the top ten and bottom ten slices) spatial overlap between the subject mask and group mask greater than 80%. The final subject pool included 315 HC and 193 SZ (*n* = 508).

### 2.2 Preprocessing

Preprocessing was performed primarily within the MATLAB software environment using Statistical Parametric Mapping (SPM12; http://www.fil.ion.ucl.ac.uk/spm/) and the FMRIB Software Library (FSL v6.0; https://fsl.fmrib.ox.ac.uk/fsl/fslwiki). Preprocessing steps included 1) rigid body motion and slice timing correction, 2) nonlinear warping to Montreal Neurological Institute (MNI) 152 coordinate space, 3) spatial resampling to 3 mm isotropic voxel spacing, 4) spatial smoothing with a 6 mm full width at half maximum (FWHM) Gaussian kernel, 5) head motion regression, detrending, despiking, low pass filtering, 6) temporal resampling to TR = 2000 ms, and finally 7) voxel time series *Z*-scoring to normalize variance.

### 2.3. Constructing Linear and Explicitly Nonlinear Whole-Brain Voxel-Wise Functional Connectivity

We construct linear as well as explicitly nonlinear global (voxel-wise) FC matrices for every subject (**Fig. 1**) (Iraji et al., 2023b). Let *X* ∈ ℝ^*n*×*v*^ be a sample of rsfMRI data where *n* is the number of time points, *v* is the number of voxels within the brain, and *x* and *y* represent any two preprocessed voxel time series such that *x, y* ∈ ℝ^1×*n*^. Thus, *x*_*i*_ is the value of voxel *x* at time point *i*. We estimate each subject’s linear whole-brain FC (LIN-wFC) as the covariance (Cov) across all pairs of brain voxels (**Eq. 1**). Because voxel time courses were *Z*-scored during preprocessing, the pairwise covariance is equal to the pairwise Pearson correlation, which is conventionally used to estimate linear FC.

**Fig. 1.**
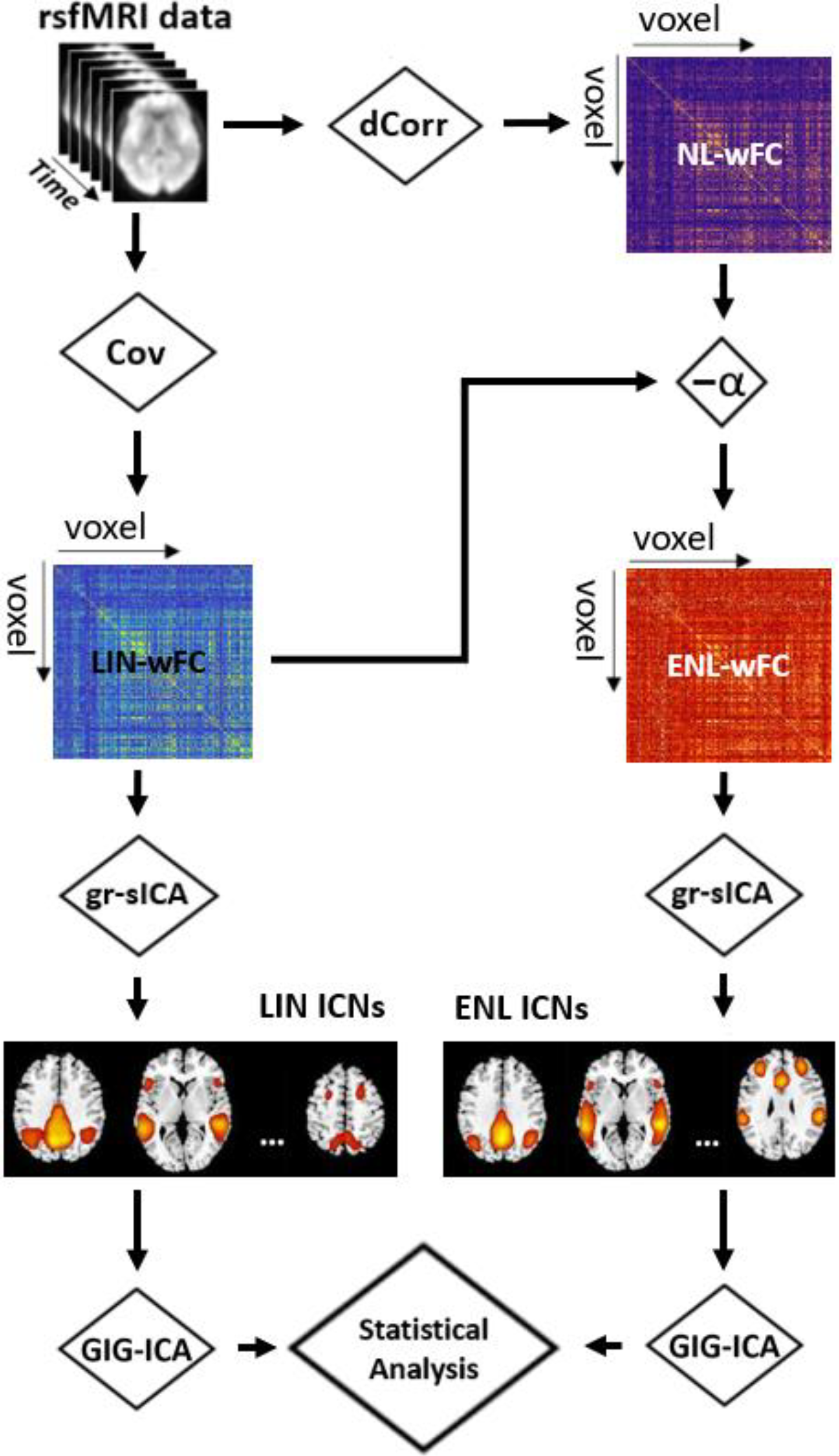
Schematic of the analysis pipeline. Preprocessed resting-state fMRI (rsfMRI) data is transformed to the connectivity domain using covariance (Cov), as a linear functional connectivity (FC) estimator, and distance correlation (dCorr), which is sensitive to both linear and nonlinear associations between voxel time series, as a nonlinear FC estimator. Explicitly nonlinear whole-brain functional connectivity (ENL-wFC) is obtained by removing the nonlinear whole-brain functional connectivity (NL-wFC) information that is linearly explained by linear whole-brain functional connectivity (LIN-wFC). Group-level spatial independent component analysis (gr-sICA) is implemented in the connectivity domain on LIN-wFC and ENL-wFC to estimate separate sets of intrinsic connectivity networks (LIN and ENL ICNs). Group information-guided ICA (GIG-ICA) is then used to estimate subject-specific ICNs, and statistical analysis is conducted on the subject-level spatial maps.

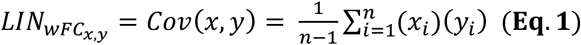

Next, we calculate the voxel-wise distance correlation (Székely et al., 2007) to construct nonlinear whole-brain FC (NL-wFC). Distance correlation is a representation of the association between random vectors based on Euclidean distances between sample observations (Székely et al., 2007) (**Eq. 2**).

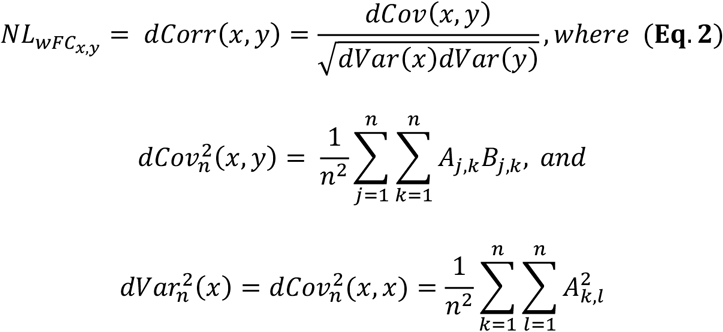

The squared sample distance covariance (dCov^2^) is calculated as the arithmetic average of the products *AB. A* and *B* represent the doubly centered Euclidean distance matrices of rsfMRI voxel time series *x* and *y* such that

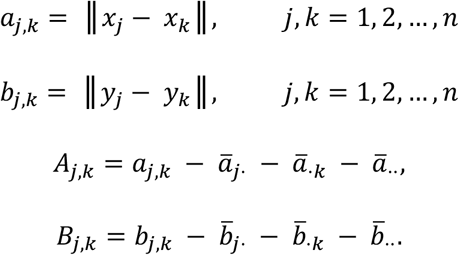

We note that distance correlation is sensitive to both linear and nonlinear dependence relations, and that the distance correlation between random vectors is zero if and only if the vectors are independent (Székely et al., 2007).

Because we are interested in analyzing the distance correlation connectivity patterns not present within Pearson correlation patterns, we remove the effect of LIN-wFC on NL-wFC using an ordinary least squares approach to estimate the explicitly nonlinear whole-brain FC (ENL-wFC) for each subject (**Eq. 3**). To estimate ENL-wFC, we first vectorize both NL-wFC and LIN-wFC. We then remove any linear relationship between NL-wFC and LIN-wFC using a regression-based method and reshape the vector of residuals into a *v* × *v* FC matrix.

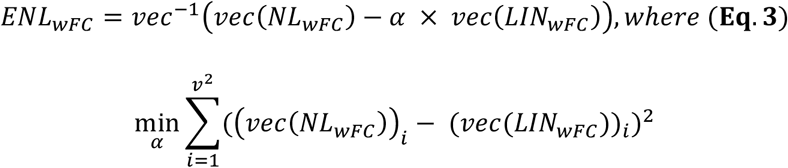

We treat the estimation of *α* as an ordinary least squares problem by finding the value of *α* which minimizes the sum of squared errors between NL-wFC and LIN-wFC. Thus, here we define the ENL-wFC for a given subject as the NL-wFC information with the linear effect of LIN-wFC removed. For each subject, the goodness of fit of the linear model was evaluated via the coefficient of determination (*R*^2^). To assess the difference in *R*^2^ between HC and SZ cohorts, we used a general linear model (GLM) to remove the effect of confounding factors including age, sex, site, and mean framewise displacement on the goodness of fit data, and we subsequently conducted a two-sided permutation test with 5000 random permutations (Krol, 2023; https://github.com/lrkrol/permutationTest).

### 2.4. Extracting Intrinsic Connectivity Networks (ICNs)

We used the Group ICA of fMRI Toolbox (GIFT v4.0; http://trendscenter.org/software/gift) (Iraji et al., 2021) to implement connectivity domain ICA (Iraji et al., 2016) and obtain separate sets of group-level ICNs from the LIN-wFC and ENL-wFC data. The implementation of group-level spatial independent component analysis (gr-sICA) was preceded by an initial subject-level multi-power iteration (Rachakonda et al., 2016) principal component analysis (PCA) step to reduce dimensionality and denoise the data (Erhardt et al. 2011). The 30 principal components (PCs) that explained the maximum variance of each subject’s respective LIN-wFC and ENL-wFC were retained for further analysis. Subject-level PCs for each FC estimator were concatenated across the component dimension, and a group-level PCA step was applied to further reduce the dimensionality of the data and decrease the computational demands of gr-sICA (Calhoun et al., 2009). The 20 group-level PCs that explained the maximum variance of each estimator-specific data set were used as the input for gr-sICA. We selected a gr-sICA model order of 20 to obtain large-scale ICNs (Iraji et al., 2016; Ray et al., 2013). To ensure the reliability of our results, ICA was implemented via the Infomax optimization algorithm (Bell & Sejnowski, 1995) 100 times with both random initialization and bootstrapping, and the most stable run was selected for further analysis. We evaluated the reliability and quality of ENL and LIN components using the ICASSO quality index (IQ), which quantifies component stability across runs (Himberg et al., 2004). To assess the difference in stability between ENL and LIN components, we conducted a two-sided permutation test with 5000 random permutations on the IQ data. Assessing component reliability was a necessary step, as previous work demonstrates that certain components may be inconsistently extracted from the data of interest (Himberg et al., 2004). In the context of fMRI ICN estimation, ICASSO IQ is often used to differentiate reliable components from components that are unstable and unfit for further analysis (Iraji et al., 2019b). A component was identified as an ICN if and only if 1) it exhibited an ICASSO IQ value exceeding .80, 2) it exhibited high visual overlap with gray matter, 3) it exhibited peak weight within gray matter, and 4) it exhibited low visual similarity to motion, ventricular, and other known artifacts. To find spatially corresponding ICNs, the spatial correlation value was computed between every pair of extracted LIN and ENL components, and components were matched in a greedy fashion. ICNs matched with a spatial correlation value exceeding .80 were classified as common (Iraji et al., 2023a) and were labeled based on their neuroanatomical distributions and the identification of ICNs from previous studies (Iraji et al., 2016). ICNs exhibiting maximum spatial correlation less than .40 were classified as unique. We used the Group ICA of fMRI Toolbox (GIFT v4.0) to implement group information-guided ICA (GIG-ICA) (Du & Fan, 2013) and reconstruct subject-specific ICNs from subject-level PCs using the group-level spatial references.

### 2.5. Assessment of Spatial Variation Among Corresponding ICNs

To assess differences in spatial variation between matched ICNs, we conducted voxel-wise paired samples two-sided *t*-tests on their *Z*-scored subject-level estimates. For a given matched ICN pair, statistical comparisons were masked for voxels exceeding *Z* = 1.96 (*p* = .05) in either group-level map (LIN or ENL), and the False Discovery Rate (FDR) method was used to correct for multiple comparisons (*q* < .05) (Benjamini & Hochberg, 1995). The automated anatomical labeling atlas 3 (AAL3) (Rolls et al., 2020) was used to localize clusters of significant voxels to anatomically defined brain regions.

### 2.6. Assessment of ICN Differences Between HC and SZ

To assess ICN differences between HC and SZ, we conducted voxel-wise independent samples *t*-tests between the estimates of common and unique ICNs derived from each cohort. We first used a GLM to remove the effect of confounding factors including age, sex, site, and mean framewise displacement on *Z*-scored subject-level ICN estimates. Voxel-wise independent samples two-sided *t*-tests were then conducted on the residual spatial maps derived from the HC and SZ groups. Common ICN statistical comparisons were masked for voxels exceeding *Z* = 1.96 (*p* = .05) in either of the group-level maps (LIN or ENL), while unique ICN comparisons were masked for voxels exceeding the same threshold in the unique group-level map. The FDR method (Benjamini & Hochberg, 1995) was used to correct for multiple comparisons (*q* < .05). The AAL3 atlas (Rolls et al., 2020) was used to localize clusters of significant voxels to anatomically defined brain regions. McNemar’s test or an exact binomial test (for *n* < 25) was used to assess the difference in statistical sensitivity between ENL and LIN estimates for every common ICN as well as across all common comparisons.

## 3. Results

### 3.1. Goodness of Fit

Goodness of fit statistics (*R*^2^) for linear regression of LIN-wFC on NL-wFC were mean ± standard deviation = .5337 ± .2009; maximum − minimum = .9413 − .0173. This indicates that, on average, much of the NL-wFC variance is captured by a linear fit. *R*^2^ is significantly higher for HC vs. SZ (*p* < .001, observed difference = .1203, Hedges’s *g* = 0.6971). The observed Hedges’s *g* value indicates the presence of a medium to large effect size.

### 3.2. Component Estimation Reliability is Greater for ENL-wFC vs. LIN-wFC

Components estimated from ENL-wFC exhibit significantly higher estimation reliability (ICASSO IQ) compared to components estimated from LIN-wFC (*p* < .001, observed difference = .037, Hedges’s *g* = 0.6441). The observed Hedges’s *g* value indicates the presence of a medium to large effect size. ENL stability indices were mean ± standard deviation = .9694 ± .0057; maximum − minimum = .9800 − .9579. LIN stability indices were mean ± standard deviation = .9324 ± .0810; maximum − minimum = .9770 − .6186.

### 3.3. Common and Unique ICNs are Identified from ENL-wFC and LIN-wFC

Within our 20-model-order gr-sICA framework, 13 ENL ICNs and 14 LIN ICNs were identified (**Fig. 2**). Among the identified ICNs, 10 exhibited maximum spatial similarity values exceeding 0.80 between their ENL and LIN estimates. These ICNs were classified as common to both ENL-wFC and LIN-wFC based on the defined criterion (Section 2.4.). Among the remaining ICNs, 2 ENL ICNs and 3 LIN ICNs exhibited maximum spatial similarity values between .40 and .80. Although several of the aforementioned ICNs attained relatively high maximum spatial similarity, we noticed distinct intensity differences across their neuroanatomical distributions which prevented common classification and labeling. Furthermore, our analysis uncovered a LIN ICN and an ENL ICN exhibiting maximum spatial similarity less than .40. These ICNs were classified as unique based on our uniqueness criterion (Section 2.4.).

**Fig. 2.**
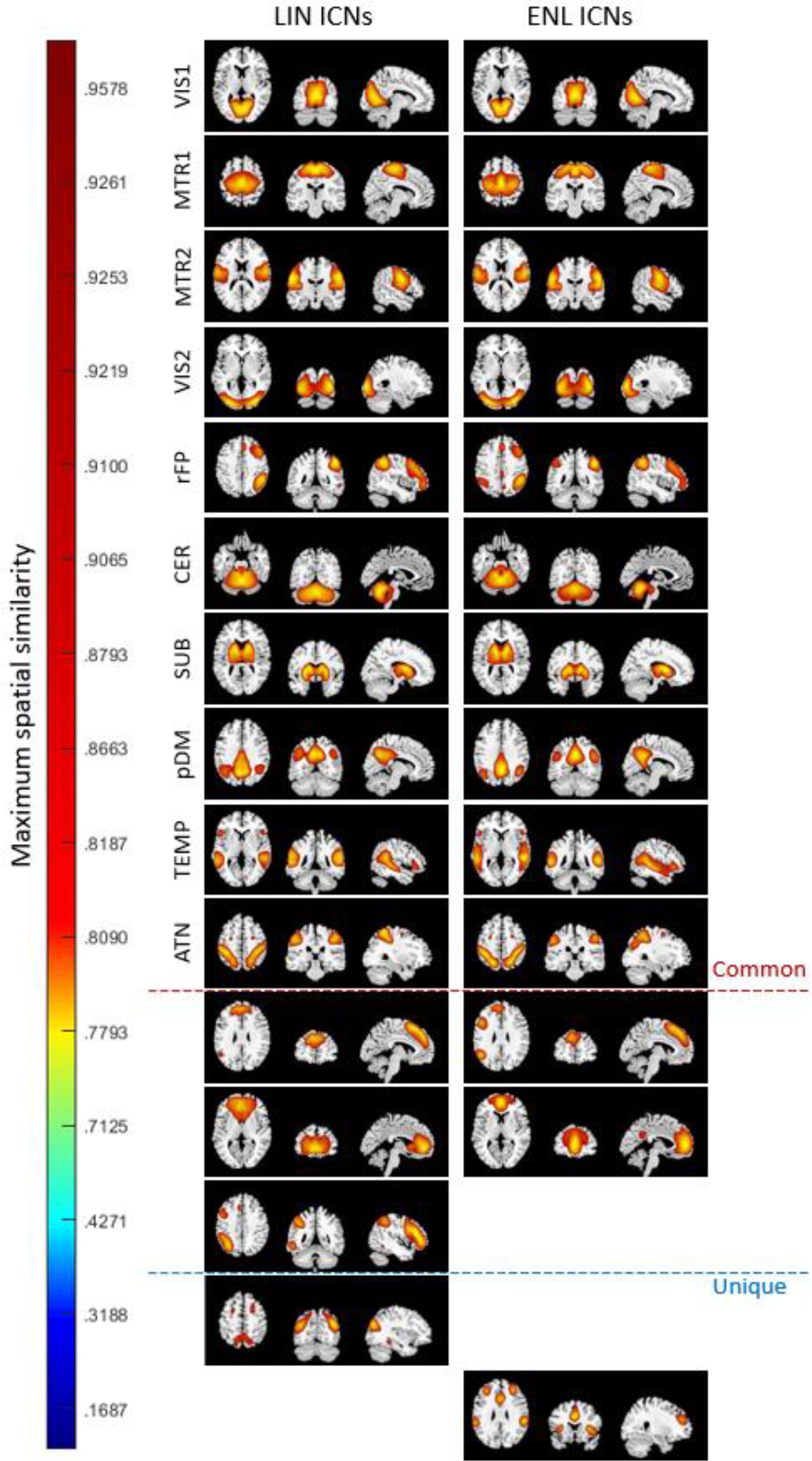
Intrinsic connectivity networks (ICNs) obtained from linear whole-brain functional connectivity (LIN-wFC) and explicitly nonlinear whole-brain functional connectivity (ENL-wFC) group-level spatial independent component analysis (gr-sICA) in the connectivity domain. ICNs are displayed thresholded at *Z* = 1.96 (*p* = .05) on the ch2bet template in order of maximum spatial similarity. Common ICNs (maximum similarity > .80) include primary visual (VIS1), primary sensorimotor (MTR1), secondary sensorimotor (MTR2), secondary visual (VIS2), right frontoparietal (rFP), cerebellum (CER), subcortical (SUB), posterior default mode (pDM), temporal (TEMP), and dorsal attention (ATN). ICNs exhibiting maximum similarity between .40 - .80 and unique ICNs (maximum similarity < .40) are also displayed.

### 3.4. ENL ICN Uniqueness: Validation

Although the spatial distribution of a component extracted from LIN-wFC gr-sICA showed similarity of .8933 to the ENL ICN that we classified as unique (**Fig. 2**), the LIN component in question was not reliably estimated, exhibiting an ICASSO quality index (IQ) value of .6186, which fell below our ICN estimation reliability threshold (.80) (Iraji et al., 2019b). Previous research supports the view that this finding strongly suggests the component is inconsistently extracted from the LIN-wFC data and unfit to be analyzed as a LIN ICN despite the similarity of its spatial distribution (Himberg et al., 2004; Iraji et al., 2019b). To validate this result, we conducted 100 additional iterations of gr-sICA on the LIN-wFC and ENL-wFC data. For every additional iteration, a randomized subset of subjects comprising 80% of the total subject pool was selected for analysis. Gr-sICA parameters were identical to those of the full analysis except for the number of Infomax runs, which was equal to five. After gr-sICA, the components extracted from each iteration were matched based on their spatial correlation values with ENL components extracted from the full analysis. Using a spatial similarity threshold of .80 and the ICN inclusion criteria specified in Section 2.4., we determined that the ICN of interest was identified in 78/100 of the additional ENL-wFC analyses (**Fig. 3A**), while it was identified in only 9/100 of the additional LIN-wFC analyses (**Fig. 3B**). This result indicates that the ICN in question cannot be reliably estimated from LIN-wFC data within a 20-model-order gr-sICA framework.

**Fig. 3.**
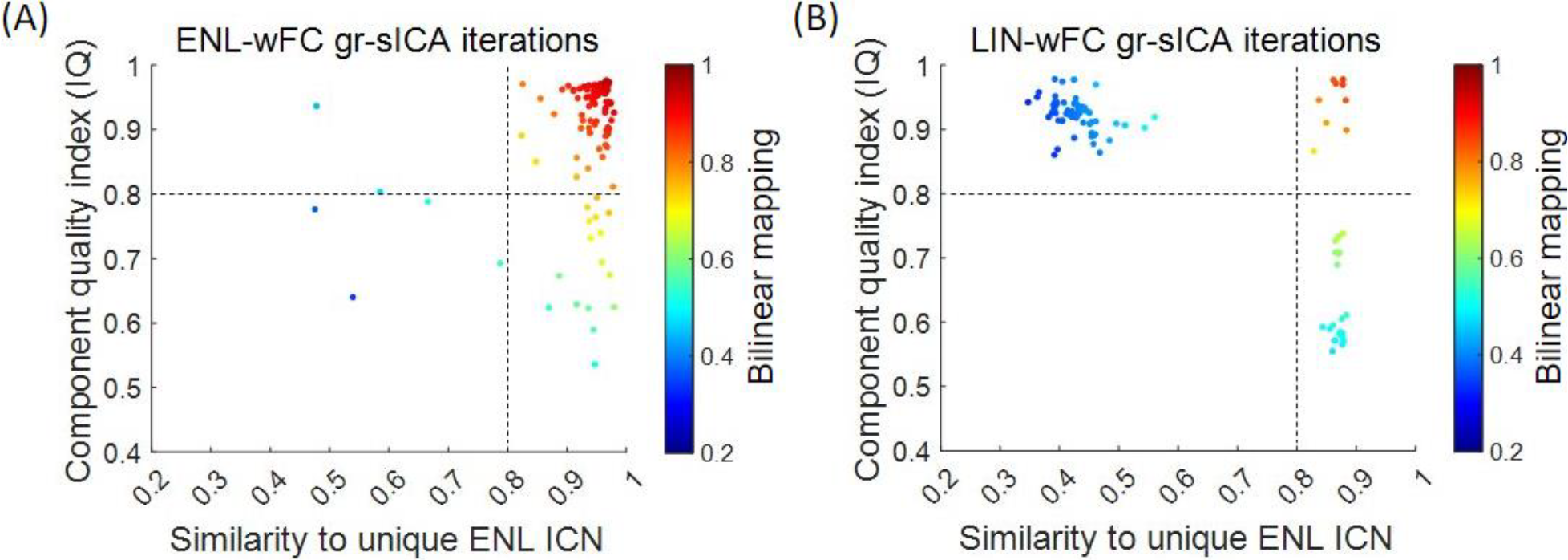
Scatterplots of nonlinear whole-brain functional connectivity (ENL-wFC) and linear whole-brain functional connectivity (LIN-wFC) group-level spatial independent component analysis (gr-sICA) iterations. Each iteration is plotted according to a colormap reflecting the bilinear mapping between the spatial correlation of the component matched with the unique ENL ICN and that component’s ICASSO quality index (IQ) value. The broken lines demarcate spatial similarity and IQ thresholds of .80. ENL-wFC gr-sICA iterations (A) cluster within the top right quadrant of the plot, indicating that matched components extracted from ENL-wFC analyses generally exhibited suprathreshold spatial similarity and suprathreshold ICASSO IQ values. This pattern is not observed in the LIN-wFC plot (B), which reveals that most LIN-wFC gr-sICA iterations failed to identify the unique ENL ICN.

### 3.5. Corresponding ENL and LIN ICNs Exhibit Unique Spatial Patterns

Our results indicate that matched ENL and LIN ICNs exhibit distinctive spatial distributions (**Fig. 4A-J**). Gradients are visually observed in ICNs associated with both lower and higher cognitive functioning, with many spatially central and core ICN regions (defined as regions that attain higher values across the spatial distribution) exhibiting greater ENL weight. For the subcortical (SUB) ICN (**Fig. 4A**), LIN weight is significantly greater within the bilateral caudate and putamen, while ENL weight is greater within bilateral thalamus. The cerebellum (CER) (**Fig. 4B)** exhibits higher ENL values within vermis lobules I-V and higher LIN values within vermis lobules VII-IX and the bilateral cerebellar hemisphere. Among networks associated with visual (Smith et al., 2009) and auditory and linguistic (Moerel et al., 2014) functioning, we find that ENL weight is predominantly greater within spatially central regions, while LIN weight is greater within peripheral areas. For instance, the primary visual (VIS1) ICN (**Fig. 4C**) exhibits a medial-lateral ENL-LIN gradient in the bilateral cortex surrounding the calcarine fissure with greater ENL weight within the cuneus. The secondary visual (VIS2) ICN (**Fig. 4D**) shows higher ENL weight within the cuneus and higher LIN weight within the bilateral inferior and middle occipital gyri. Temporal (TEMP) ICN (**Fig. 4E**) spatial variation follows a similar center-periphery pattern, with greater ENL weight in the superior temporal gyri and greater LIN weight within the supramarginal gyri and bilateral inferior frontal triangularis.

**Fig. 4.**
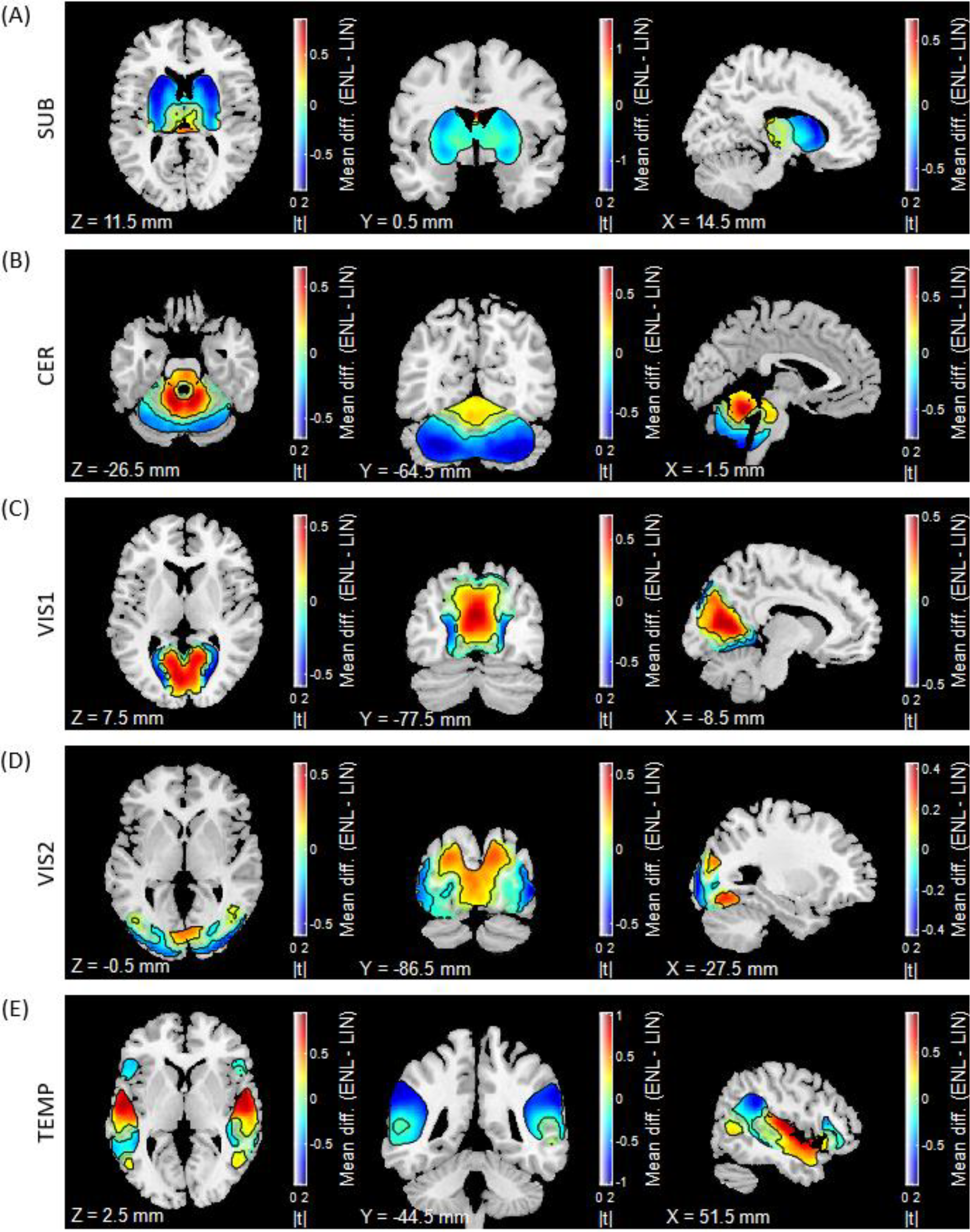
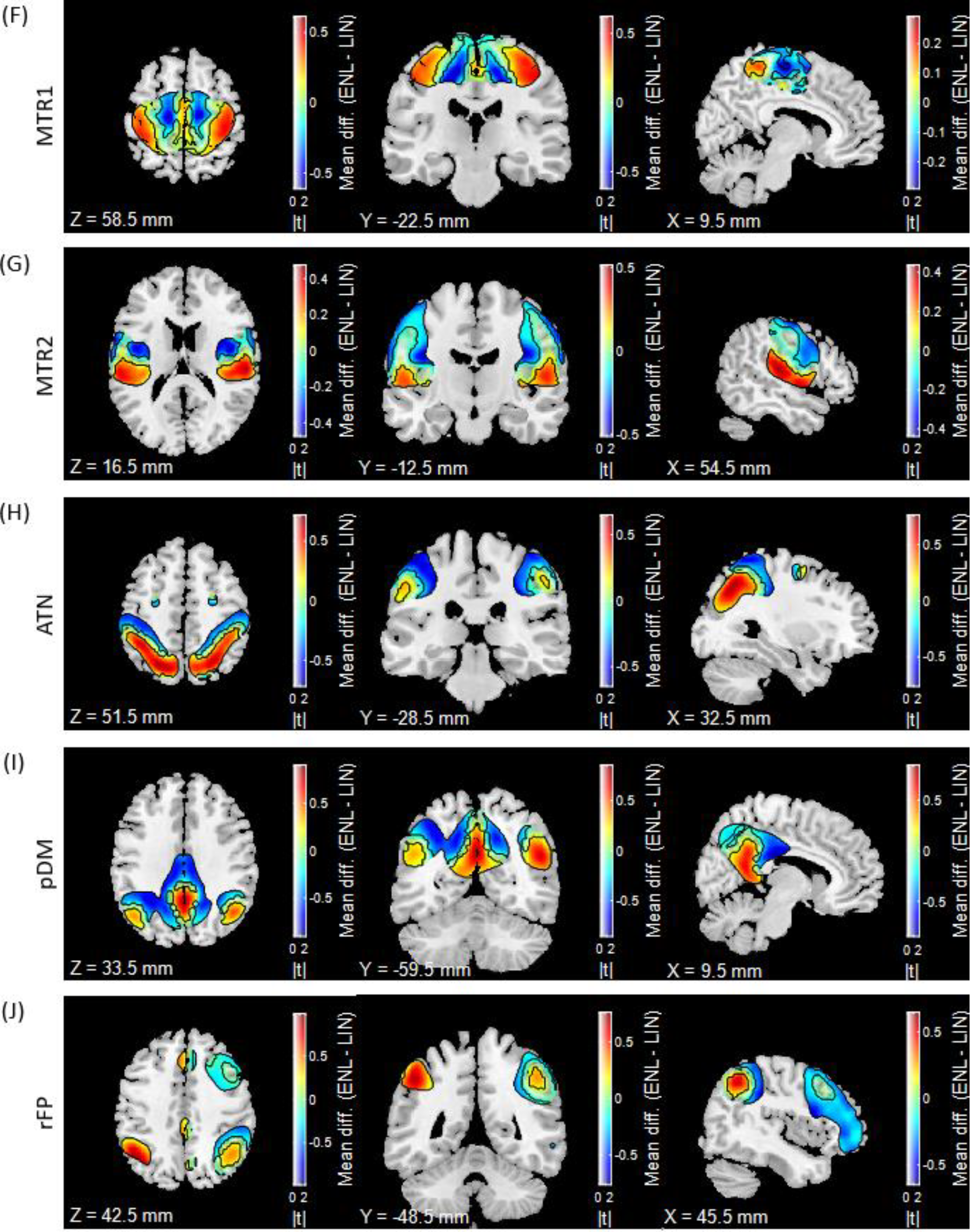
Assessment of intrinsic connectivity network (ICN) spatial variation. Warmer hues indicate ENL > LIN, while cooler hues indicate LIN > ENL. Contours indicate statistical significance (*q* < .05). Displayed ICNs include subcortical (SUB) (A), cerebellum (CER) (B), primary (VIS1) (C) and secondary (VIS2) (D) visual, temporal (TEMP) (E), primary (MTR1) (F) and secondary (MTR2) (G) sensorimotor, dorsal attention (ATN) (H), posterior default mode (pDM) (I), and right frontoparietal (rFP) (J). Results are overlaid on the ch2bet template with X, Y, and Z coordinates listed relative to the origin in Montreal Neurological Institute (MNI) 152 space. Dual code visualization was adapted from sample scripts provided by Allen et al. (2012).

Whereas both the primary and secondary sensorimotor ICNs (MTR1 and MTR2) show ENL-LIN gradients (**Fig. 4F-G**), MTR1 comparisons reveal a medial-lateral pattern between the paracentral lobules and pre- and postcentral gyri, while MTR2 comparisons reveal an inferior-superior gradient between the superior temporal lobe and pre- and postcentral gyri. ICNs associated with higher cognitive functions such as attention (Szczepanski et al., 2013), social cognition and self-referential processing (Wang et al., 2020), and executive control (Niendam et al., 2012) exhibit core-periphery gradients. The dorsal attention (ATN) (**Fig. 4H**) ICN shows higher ENL weight in the superior parietal lobules and higher LIN weight in the postcentral gyri. The posterior default mode (pDM) ICN (**Fig. 4I**) exhibits higher ENL values in the precuneus and bilateral angular gyri with higher LIN values in the middle and posterior cingulate. The right frontoparietal (rFP) ICN (**Fig. 4J**) exhibits higher ENL values within the angular gyri (particularly within the left angular gyrus) and higher LIN values within the right inferior parietal lobule, right middle frontal gyrus, and right inferior frontal triangularis.

### 3.6. ENL ICN Estimates Exhibit Enhanced Sensitivity to Differences Between HC and SZ, and Unique ENL (but Not LIN) ICN Estimates Reflect Group Differences

ENL estimates exhibit an overall greater degree of sensitivity to differences between HC and SZ compared to LIN estimates (*p* < .001) in addition to revealing a larger total number of significant voxels across all comparisons. Moreover, the ENL estimates of ICNs associated with auditory and linguistic (Bhaya-Grossman & Chang, 2022; Moerel et al., 2014; Rupp et al., 2022), sensorimotor (Caspers et al., 2021), and self-referential (Wang et al., 2020) processes exhibit enhanced sensitivity to differences between HC and SZ (**Fig. 5A-C**). For example, while both sets of comparisons detected differences within TEMP ICN regions comprising the primary auditory and auditory association cortex, ENL comparisons are more sensitive (*p* < .001), revealing clusters that are more numerous with augmented volumes and effect sizes (**Fig. 5A**). LIN and ENL testing detected higher HC values within the bilateral superior temporal gyri and temporal poles, bilateral insula, bilateral Heschl’s gyrus, bilateral Rolandic operculum, and right middle temporal gyrus along with higher SZ values within the right supramarginal gyrus. However, ENL comparisons revealed larger numbers of significant voxels across these regions. Additionally, ENL comparisons uncovered higher HC values within the left middle temporal gyrus and higher SZ values within the left supramarginal gyrus, both of which were missed for significance by LIN comparisons. We visually observe that TEMP regions showing higher values for HC exhibit higher ENL values compared to LIN (as revealed by our assessment of spatial variation; Section 3.5.) with the exception of the left middle temporal gyrus, while TEMP regions showing higher SZ values exhibit higher LIN values relative to ENL.

**Fig. 5.**
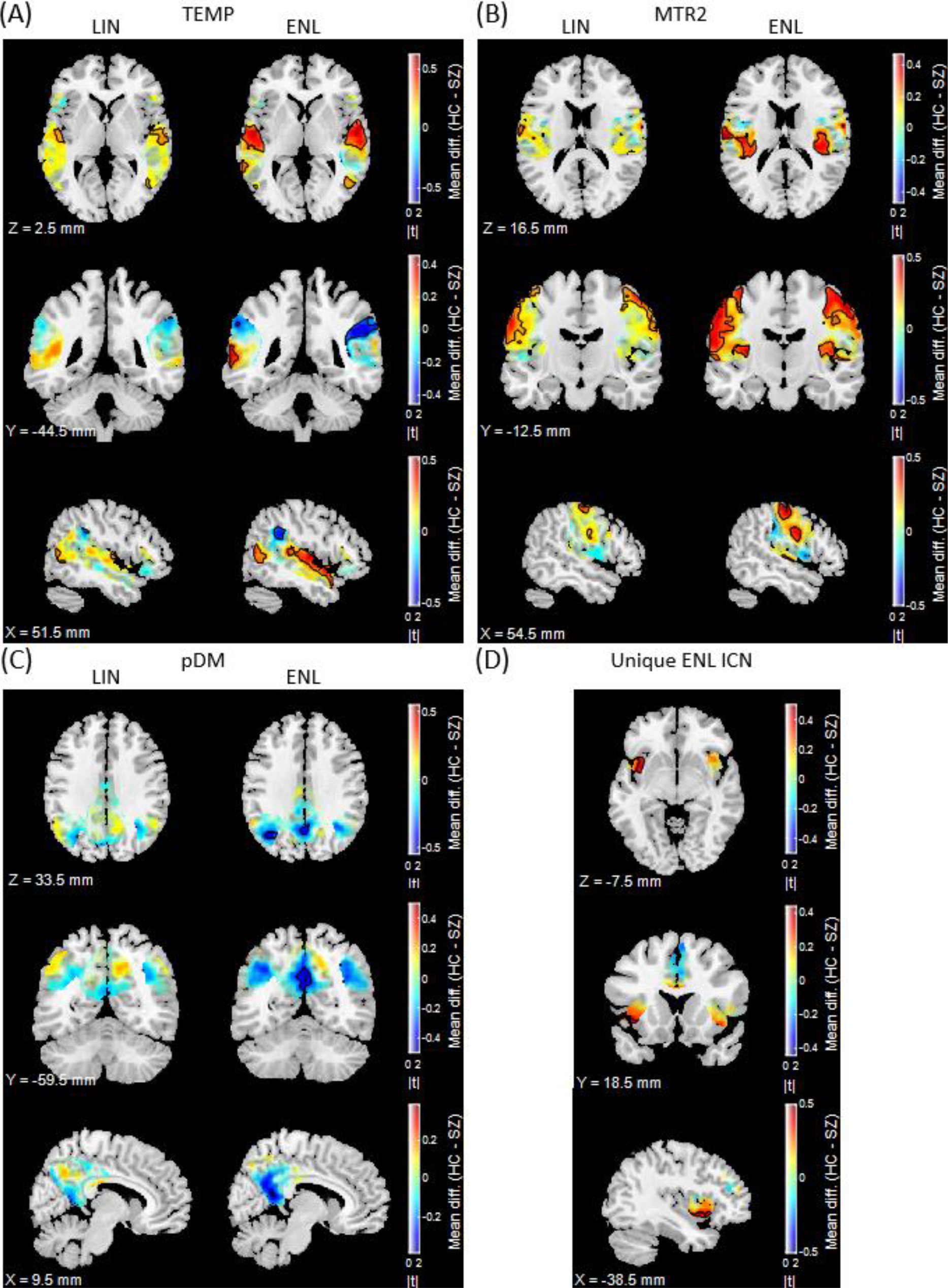
Statistical comparisons between subject-level temporal (TEMP) (A), secondary sensorimotor (MTR2) (B), posterior default mode (pDM) (C), and unique ENL (D) ICN estimates derived from distinct clinical cohorts (HC and SZ). In A-C, results from LIN comparisons are located on the left, while results from ENL comparisons are located on the right. Warmer hues indicate HC > SZ, while cooler hues indicate SZ > HC. Contours indicate statistical significance (*q* < .05). Results are overlaid on the ch2bet template with X, Y, and Z coordinates listed relative to the origin in MNI 152 space. Dual code visualization was adapted from sample scripts provided by Allen et al. (2012).

A similar pattern of higher ENL sensitivity was obtained for secondary sensorimotor (MTR2) (**Fig. 5B**) (*p* < .001) and posterior default mode (pDM) (**Fig. 5C**) (*p* < .001) comparisons. For MTR2, both sets of comparisons revealed greater MTR2 weights for HC in the bilateral postcentral gyri. However, ENL comparisons detected more extensive clusters along with the additional finding that HC exhibit greater MTR2 values within the bilateral posterior insula. For pDM estimates, ENL comparisons revealed clusters of significantly higher values for SZ within the precuneus and left angular gyrus, while LIN comparisons identified only two significant voxels. Visual inspection reveals that ENL pDM regions showing higher SZ values also exhibit higher ENL values within the pDM ENL-LIN spatial gradient (Section 3.5.). We note that LIN estimates exhibit a greater degree of statistical sensitivity for cerebellum (CER) (*p* < .001), primary visual (VIS1) (*p* < .001), secondary visual (VIS2) (*p* < .001), primary sensorimotor (MTR1) (*p* < .05), and right frontoparietal (rFP) (*p* < .001) ICNs. While unique LIN ICN comparisons failed to detect any significant group differences, we found that unique ENL ICN comparisons revealed a cluster of voxels within the left anterior insula that distinguish clinical cohorts, with HC exhibiting significantly greater values than SZ (**Fig. 5D**). Results from all clinical cohort voxel-wise statistical comparisons and sensitivity tests can be found in Supplemental Material **Table S1** and **Table S2**.

## 4. Discussion

Linear FC analysis remains a fruitful method for extracting valuable information from fMRI data. However, despite its usefulness and ease of interpretation, various brain processes also exhibit nonlinear aspects (Friston, 2001; Singer, 2013), suggesting that linear FC provides us with a limited view of the data and neurocognitive hypothesis space. While previous rsfMRI studies have identified evidence of nonlinearity and its prospective role in differentiating clinical cohorts (Morioka et al., 2020; Motlaghian et al., 2022; Motlaghian et al., 2023), here we advance a novel approach to estimate ICNs from explicitly nonlinear whole-brain FC (ENL-wFC) constructed from residual distance correlation information, demonstrating the potential of connectivity domain ICA (Iraji et al., 2016) and nonlinear information to shape the predictive clinical landscape and inform systems neuroscience theorizing.

We find that ICN estimates extracted from ENL-wFC exhibit higher reliability than those extracted from LIN-wFC (Section 3.2.), and that unique ICNs are identified from each FC estimator (Section 3.3.). Our validation analysis (Section 3.4.) supports these findings. That our approach can recover ICNs that would be missed by conventional linear FC analysis underscores the importance of bringing nonlinearity within the scope of fMRI FC research, as ICNs estimated from connectivity patterns that are not present in LIN-wFC may be altered in psychiatric conditions such as SZ.

We also find that corresponding (spatially matched) ICNs exhibit striking ENL-LIN spatial gradients (Section 3.5.). Among potential explanations for our findings, the presence of greater ENL weight within core ICN regions could be reflective of stronger signal within core areas. However, we note that this explanation does not align with the detection of differences between HC and SZ cohorts (Section 3.6.). Conceptually, the ICNs extracted from ENL-wFC represent independent data sources comprised of elements whose distance correlation values deviate from a linear relationship with Pearson correlation. Therefore, the identified gradients may reflect actual differences in the complexity of the underlying FC relationships, which would merit further investigation of their potential cognitive and clinical significance. Future work will investigate potential explanations of the observed gradients.

Furthermore, the finding that ENL estimates exhibit an overall greater degree of sensitivity to differences between HC and SZ compared to LIN demonstrates the potential of nonlinear information to play a role within predictive models of diagnosis. The ENL counterparts of specific ICNs that have been reported as disrupted in SZ including temporal (TEMP), secondary sensorimotor (MTR2), and posterior default mode (pDM) exhibit heightened sensitivity to differences between HC and SZ vs. LIN (Section 3.6.). ENL TEMP comparisons revealed larger clusters of voxels within auditory and language-related regions that have been previously associated with SZ and positive symptoms such as auditory verbal hallucinations in both tfMRI (Calhoun et al., 2012; Kim et al., 2009) and rsfMRI (Alderson-Day et al., 2015; Iraji et al., 2019b). For example, ENL testing revealed expansive clusters within the superior temporal gyri, which are known to implement acoustic-phonetic computations (Bhaya-Grossman & Chang, 2022). Notably, ENL TEMP comparisons also identified a sizable volume attaining significantly higher SZ values within the right supramarginal gyrus, which has been shown to play a role in phonological decision-making (Hartwigsen et al., 2010). By contrast, the right supramarginal gyrus was almost entirely missed for significance by LIN TEMP comparisons. We find that ENL MTR2 comparisons revealed greater numbers of significant voxels compared to LIN within sensorimotor regions previously implicated in SZ (Kaufmann et al., 2015; Iraji et al., 2019b) as well as clusters within the bilateral posterior insula hidden from LIN MTR2 comparisons. Moreover, ENL pDM testing revealed SZ hyperconnectivity within the precuneus and left angular gyrus that were missed by LIN analysis, which are core regions of the pDM that have been associated with reflective, internally focused cognitive processes thought to be relevant to SZ diagnosis and symptoms (Garrity et al., 2007). Additionally, unique LIN ICN comparisons failed to distinguish clinical cohorts while unique ENL ICN comparisons identified higher values for HC within the left anterior insula, which is characteristically associated with an ICN involved in event and stimulus salience processing known to be compromised in SZ (Palaniyappan & Liddle, 2012). Overall, our results demonstrate that nonlinear statistical dependencies in fMRI data can be leveraged to distinguish clinical cohorts and warrant further investigation of the relationship between features extracted from measures that are sensitive to nonlinearity and the presentation of psychosis.

While our previous work proposed this conceptual framework (Iraji et al., 2023b), here we advance and rigorously investigate the framework by providing an in-depth quantitative analysis of ENL and LIN ICNs, their spatial variation, and their sensitivity to differences between HC and SZ cohorts. However, the current analysis has methodological and interpretive limitations that are important to recognize. First, we note that alternative models of the relationship between NL-wFC and LIN-wFC can be leveraged when estimating ENL-wFC. Therefore, we do not claim that the current method of estimation is decisive or definitive to the potential exclusion of methods designed to estimate ENL-wFC using alternative models. Future work will investigate the use of alternative models with the aim of providing increasingly robust and precise characterizations of whole-brain connectivity features not present within linear connectivity patterns. Second, we note that while our approach may share certain conceptual similarities with methods that construct nonlinear fMRI connectivity using features derived from time series (pairwise) relationships (Motlaghian et al., 2022; Motlaghian et al., 2023), we do not necessarily expect the findings of these distinct approaches to converge due to substantial differences in methodology (Section 1.). Thus, we leave any speculation about the relationship between features extracted from these methods as an open empirical question for future investigation. Finally, while our results warrant further investigation into the potential neurocognitive and psychiatric roles of ENL ICNs, we maintain that going beyond association will likely require developing interventions that can effectively tie the extracted features to the causal outcomes of cognitive operations, psychiatric diagnosis, and symptoms.

Ongoing and future work will also focus on replicating our results in large-scale B-SNIP transdiagnostic rsfMRI data sets (Meda et al., 2012; Meda et al., 2015), on utilizing ENL ICNs to distinguish a broader array of clinical cohorts, on analyzing associations with cognitive and symptom scores, and on analyzing the temporal (Iraji et al., 2021) and spatial (Bhinge et al., 2019; Iraji et al., 2020; Long et al., 2021) dynamics exhibited by ENL ICNs during task performance and at rest.

## 5. Conclusion

Here, we advance a novel approach to estimate ICNs from explicitly nonlinear whole-brain FC (ENL-wFC). We demonstrate that our approach reveals unique spatial variation within the ENL estimates of ICNs identified within the existing literature and that spatially common as well as unique ENL ICNs thought to be relevant to SZ and its symptoms exhibit heightened sensitivity to differences between HC and SZ cohorts. In summary, our research emphasizes the importance of bringing nonlinearity within the aperture of fMRI FC analysis and the power of connectivity domain ICA to transform predictive clinical and systems neuroscience research.

## Funding acknowledgement

This work was supported by the National Institutes of Health (NIH) grant number R01 5R01MH119251 to Armin Iraji, National Science Foundation (NSF) grant number 2112455 to Vince Calhoun, Department of Veterans Affairs Senior Research Career Scientist Award number 1IK6CX002519 to Judith Ford, and the National Center for Research Resources at the National Institutes of Health grant numbers NIH 1 U24 RR021992 (Function Biomedical Informatics Research Network) and NIH 1 U24 RR025736-01 (Biomedical Informatics Research Network Coordinating Center; http://www.birncommunity.org). Pablo Andrés Camazón has received grant support from Programa Intramural de Impulso a la I+D+i 2023 (Instituto de Investigación Sanitaria Gregorio Marañón).

