## Supplemental Material for "Networks extracted from nonlinear fMRI connectivity exhibit unique spatial variation and enhanced sensitivity to differences between individuals with schizophrenia and controls"

**Table S1**. Results from voxel-wise statistical comparisons between explicitly nonlinear (ENL) and linear (LIN) intrinsic connectivity network (ICN) estimates derived from healthy controls (HC) and individuals with schizophrenia (SZ).

| ICN | Number of significant voxels (ENL) | Number of significant voxels (LIN) |
|---|---|---|
| SUB | 131 | 141 |
| CER | 424 | 1184 |
| VIS1 | 361 | 558 |
| VIS2 | 12 | 405 |
| TEMP | 1236 | 306 |
| MTR1 | 59 | 84 |
| MTR2 | 1513 | 680 |
| ATN | 0 | 0 |
| pDM | 129 | 2 |
| rFP | 0 | 12 |
| Unique LIN | - | 0 |
| Unique ENL | 60 | - |
| **Total** | 3925 | 3372 |

**Table S2**. Results from statistical sensitivity testing between explicitly nonlinear (ENL) and linear (LIN) intrinsic connectivity network (ICN) estimates derived from healthy controls (HC) and individuals with schizophrenia (SZ). For ENL vs. LIN, greater statistical sensitivity is indicated at the following significance levels: ***$p$ < .001, **$p$ < .01, *$p$ < .05. Odds ratio (OR) is used as an indicator of effect size.

| ICN | Number of significant voxels identified by ENL but not LIN | Number of significant voxels identified by LIN but not ENL | Effect size (OR) |
|---|---|---|---|
| SUB | 101 | 111 | - |
| CER | 256 | 1006*** | 3.93 |
| VIS1 | 154 | 351*** | 2.28 |
| VIS2 | 1 | 394*** | 394 |
| TEMP | 973*** | 43 | 22.63 |
| MTR1 | 33 | 58* | 1.76 |
| MTR2 | 959*** | 126 | 7.61 |
| ATN | 0 | 0 | - |
| pDM | 128*** | 1 | 128 |
| rFP | 0 | 12*** | Undefined |
| **Total** | 2605*** | 2102 | 1.24 |
